# Event-triggered STED imaging

**DOI:** 10.1101/2021.10.26.465907

**Authors:** Jonatan Alvelid, Martina Damenti, Ilaria Testa

**Affiliations:** Department of Applied Physics and Science for Life Laboratory, KTH Royal Institute of Technology, 100 44 Stockholm, Sweden

**Author notes:** Corresponding author: Ilaria Testa, **Email:**.

## Abstract

The observation of protein organization during cellular signalling calls for imaging methods with increased spatial and temporal resolution. STED nanoscopy can access dynamics of nanoscale structures in living cells. However, the available number of recordable frames is often limited by photo-bleaching. Here, we present an automated method, event-triggered STED, which instantly (< 40 ms) images synaptic proteins with high spatial and temporal resolution (~30 nm, 2.5 Hz) in small regions upon and at the site of local calcium sensing.

## Main

STED nanoscopy has been successfully used to image a variety of structures in both living cells and tissues even dynamically^1^. The temporal resolution of STED nanoscopy usually depends on the size of the region of interest (ROI) to be imaged, owing to its most common point-scanning implementation. This means that the technique can achieve high frame rate imaging (1–30 Hz) in small regions of interest (1–5 μm^2^)^2–4^, while for imaging larger fields of view, up to 80 x 80 μm^2^, it takes up to minutes to acquire a single frame^5^. This sets the current trade-off between spatial and temporal resolution where fast dynamics inside cells can only be followed in sufficiently small areas that are often difficult to pin-point due to the loss of the larger cellular context. Parallelized STED methods^6, 7^ have tried to overcome this trade-off by minimizing the number of scanning steps during imaging, however they are currently limited by the camera frame rate and depletion power. Moreover, while STED nanoscopy is capable of a high temporal resolution, it is also susceptible to photobleaching and photodamage, which limits the total number of recordable frames^8, 9^. Sample-adaptive scanning approaches^10, 11^ have helped in minimizing the light dose and the recording time during imaging. However, they have not yet been synchronized to and triggered by real-time changes in the sample such as intensity spikes, local movement, or morphological changes, neither have they been automated to switch between distinct imaging modalities. Approaches often called intelligent microscopy attempts to provide solutions, but they so far have focused on increasing the throughput of large-scale and relatively slow events such as cellular division with comparatively low resolution imaging methods^12^. Implementations incorporating nanoscopy methods and operating at a millisecond time scale to investigate nanoscale details of fast dynamic processes such as sensing are so far missing. Therefore, it is difficult with current STED microscopes to observe dynamic events efficiently and rapidly in cells, either because they are hard to find, they happen too fast, or because the availability of frames before bleaching is not high enough. However, if the user would know where and when to image the cellular process of interest, the quality, throughput, speed, and length of the observation would increase substantially and allow to unravel dynamics of the process comprehensively.

Here, we present a novel sample-adaptive microscopy method called event-triggered STED (etSTED), which enables rapid STED nanoscopy acquisitions upon and at the site of automatically detected local calcium sensing. It does so by combining fast (20 Hz) widefield imaging, which facilitate the detection and localization of sensing events, with STED imaging, for high-resolution acquisition at the site of a detected event within a temporal window of 40 ms. The method runs a real-time analysis pipeline, flexibly optimized to the problem at hand, on every recorded widefield frame. In our implementation, the analysis pipeline detects sensing events such as localized calcium spikes in neurons, which triggers STED timelapse imaging of actin, tubulin, or proteins within synaptic vesicles in the very same area. Thanks to the generalized implementation of etSTED any other combination of dynamic events and structures can be observed. Furthermore, the adaptive method does not lend itself limited to use specifically with widefield and STED imaging, but instead the concept is applicable to other combinations of fast and high-resolution imaging methods.

## Results

Our automated etSTED acquisition scheme can trigger high-resolution STED imaging upon detected events in widefield images in the millisecond-regime (Fig. 1a–b). The widefield images are processed with an optimized analysis pipeline, which returns a set of coordinates of any detected event. If an event, such as an intensity variation, is detected, the widefield imaging is stopped and the STED acquisition started in an area around the detected coordinate with pre-determined image acquisition parameters. When the STED image or timelapse is acquired, all the data regarding the event is saved. This includes a widefield timelapse leading up to the detected event, the scanned STED image or timelapse, and a log file summarizing the parameters and timings of the event detection and scanning. The saved auxiliary data is important to confirm the validity of the event in a post-acquisition analysis and allow quantification of the temporal and larger spatial context of the event. The microscope then returns to the previous settings and another continuous widefield recording is instantly initiated. The method can run indefinitely, and the focus lock in place in the microscope^5^ (Supplementary Fig. 2) maintains a stable sample throughout the experiment and ensures that the widefield and STED images are recorded in the same focal plane.

**Figure 1.**
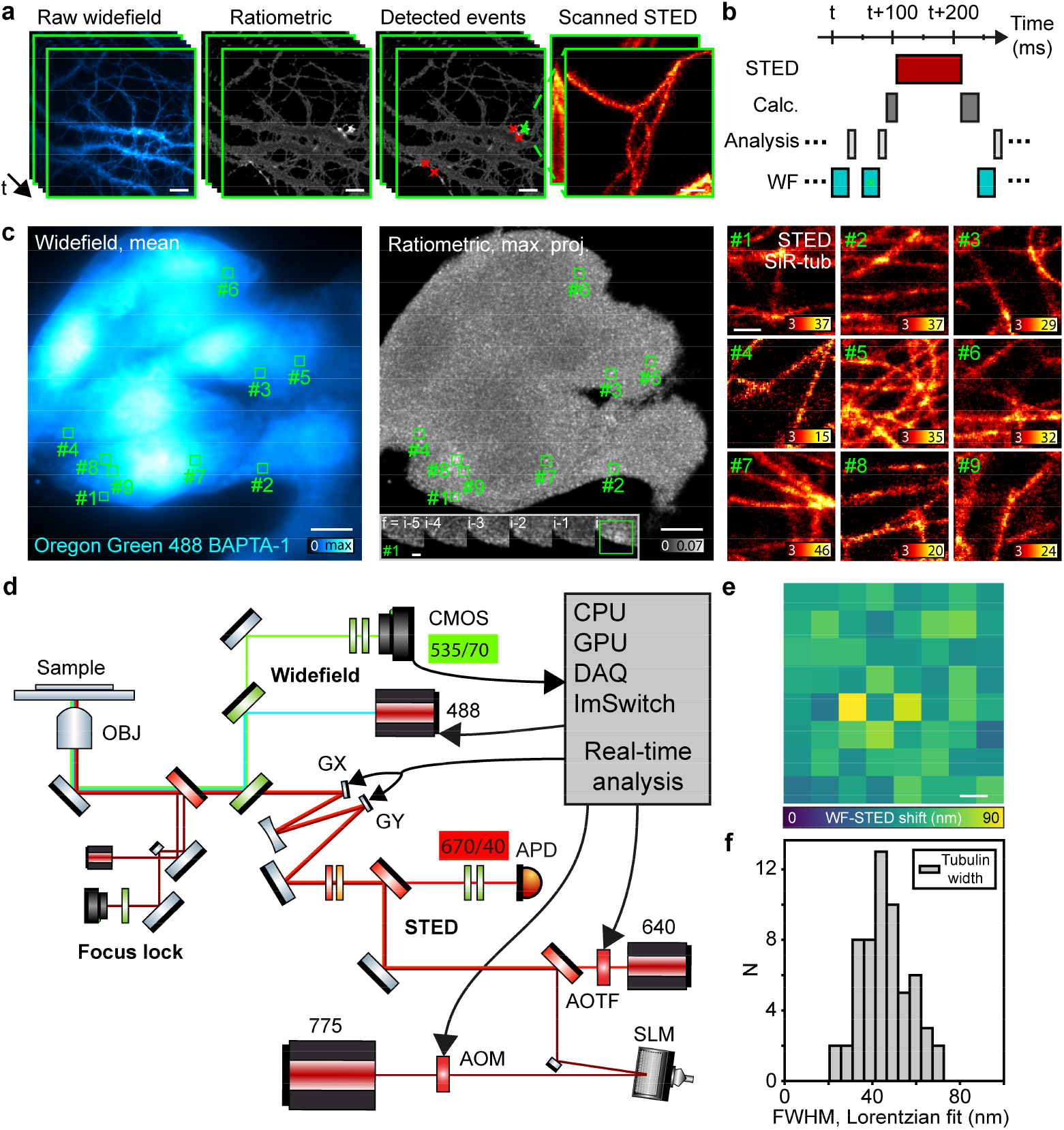
Overview of event-triggered STED. **a**, Scheme of etSTED with widefield calcium imaging of Oregon Green 488 BAPTA-1 in neurons, corresponding ratiometric images, detected coordinates of events, and locally scanned etSTED images of SiR-tubulin. Green cross indicates the ratiometrically brightest detected event that triggered STED imaging, red crosses indicate additionally detected events in the same widefield frame. **b**, Timeline of typical etSTED experiment, with widefield, analysis pipeline, overhead, and STED imaging time. **c**, etSTED experiment in HeLa cells with BAPTA-1 calcium imaging with mean widefield (left) and maximum projection ratiometric (center) images, showing the location of 9 detected events throughout the experiment, and 9 event-triggered 2 x 2 μm^2^ STED images (right). Inset (center) shows a zoomed-in view of the ratiometric image around event #1 for the five frames leading up to the event. **d**, Schematic view of the microscope setup, combining STED and widefield imaging with real-time analysis and acquisition control in ImSwitch that controls laser powers, detectors, and scanning galvanometric mirrors. **e**, Characterization of residual shift after coordinate transformation between widefield and scanning space across the FOV. N = 1048 beads. **f**, Characterization of the achieved resolution in etSTED images across multiple experiments with SiR-tubulin in HeLa cells and across the FOV, as fitted FWHM in line profiles across single microtubules. N = 69 events in N = 8 cells. Scale bars, 10 μm (**a**,**c** widefield, ratiometric, **a** detected events, **e**), 2 μm (**c** inset) and 500 nm (**a**,**c** STED).

In this work an optimized analysis pipeline for detecting calcium intensity spikes in BAPTA-1-labelled samples is used (Supplementary Fig. 1, Supplementary Note 1), but the software and method lends themselves to being used with any event-detection pipeline. A comparison between the current and previous widefield frame (Fig. 1c, left) gives a map of ratiometric intensity changes in the sample (Fig. 1c, middle). A peak-detection analysis on the ratiometric image can then extract coordinates relating to events of intensity changes in the widefield image (Fig. 1c, green boxes). If more than one event is detected, the ratiometrically brightest is chosen. At the site of a detected event, STED imaging is subsequently performed, as in the examples with HeLa cells and SiR-tubulin (Fig. 1c, right). An 80 x 80 μm^2^ region of the sample is surveyed at any time (Fig. 1c, left), and STED imaging can take place at any position. The custom-built setup (Fig. 1d, Supplementary Fig. 2) combines STED with widefield imaging by spectrally separating the two techniques and is controlled with the open-source software ImSwitch^13^. The control software integrates all hardware, for both STED imaging and widefield imaging, allowing fast control of the integrated components, ultimately enabling the instant event-triggered method that requires rapid feedback between the two modalities and their hardware components: lasers, scanners, cameras, and acousto-optic modulators and tuneable filters. The etSTED method is fully controlled via a widget in ImSwitch. We further validated that a third-order polynomial coordinate transformation^14^ between the widefield space and scanning space gives accurate transformations across the field of view (FOV) (Fig. 1e). The transformation is calibrated by detecting fluorescent beads in the two imaging modalities and fitting the coefficients in the polynomial transformation. The accuracy was analysed by transforming widefield coordinates of all detected beads into scanning space and comparing the transformed coordinates with those of the detected beads in the scanned image. The distance between coordinate pairs, transformed and detected, was calculated to generate a map of the accuracy which shows no spatial correlation and a mean transformation error of 54 nm across the FOV, which can be compared to the widefield pixel size of 100 nm. The spatial resolution in the calcium-triggered STED images across the FOV was quantified by measuring the widths of microtubules, resulting in a distribution of 46 ± 11 nm (Fig 1f), confirming that the STED image resolution and quality is not affected by the triggering method.

We applied the method to study the rearrangement of synaptic proteins at high spatio-temporal resolution during calcium activity in hippocampal neurons (Fig. 2). The authenticity and efficiency of the detection of calcium spikes in neurons was investigated (Fig. 2a–c). By applying the analysis pipeline post-acquisition on a widefield timelapse recording (Fig. 2a) we could extract calcium intensity traces at the sites of detected events also after the detection (Fig. 2b, left). We can confirm that the detected events are, on average, in 91% (0.91 ± 0.19) of the cases true calcium spike events (Fig. 2c), as the characteristic initial spike and decay across hundreds of milliseconds can be seen. Moreover, in over half of the analysed timelapses 100% of the detected events were true. A few of the detected events in the timelapses do not show a real spike at t = 0, and through manual annotation of the timelapses it is further confirmed that these are false detections, and in most cases are due to rapid movement of small dendritic filopodia-like structures in neuronal cells. Important to note is that these events can be sorted out in post-acquisition analysis by inspecting the calcium trace up to the point of event detection (Supplementary Fig. 3, Supplementary Note 2). The detected movement suggests another potential application of the method whereby detection of cellular movement can be used to trigger STED imaging. Furthermore, by manually annotating calcium events in the timelapse recordings we can see which of the events are detected with the analysis pipeline (Fig. 2b, middle). On average, 76% (0.76 ± 0.16) of the events in the neurons are detected by the analysis pipeline (Fig 2c), and only a few are missed, with generally a lower ratiometric peak value that have fallen below the applied ratiometric threshold. This means that the true positive event detection ratio (0.91) is higher than the detected events ratio (0.76), which is due to the pipeline being optimized for accuracy to minimize false detections, but regardless detects most of the events. Lastly, to further confirm the detect events as calcium spikes we extract calcium traces from random positions inside the neurons in the very same timelapses (Fig. 2b, right), which show expected flat calcium signal profiles.

**Figure 2.**
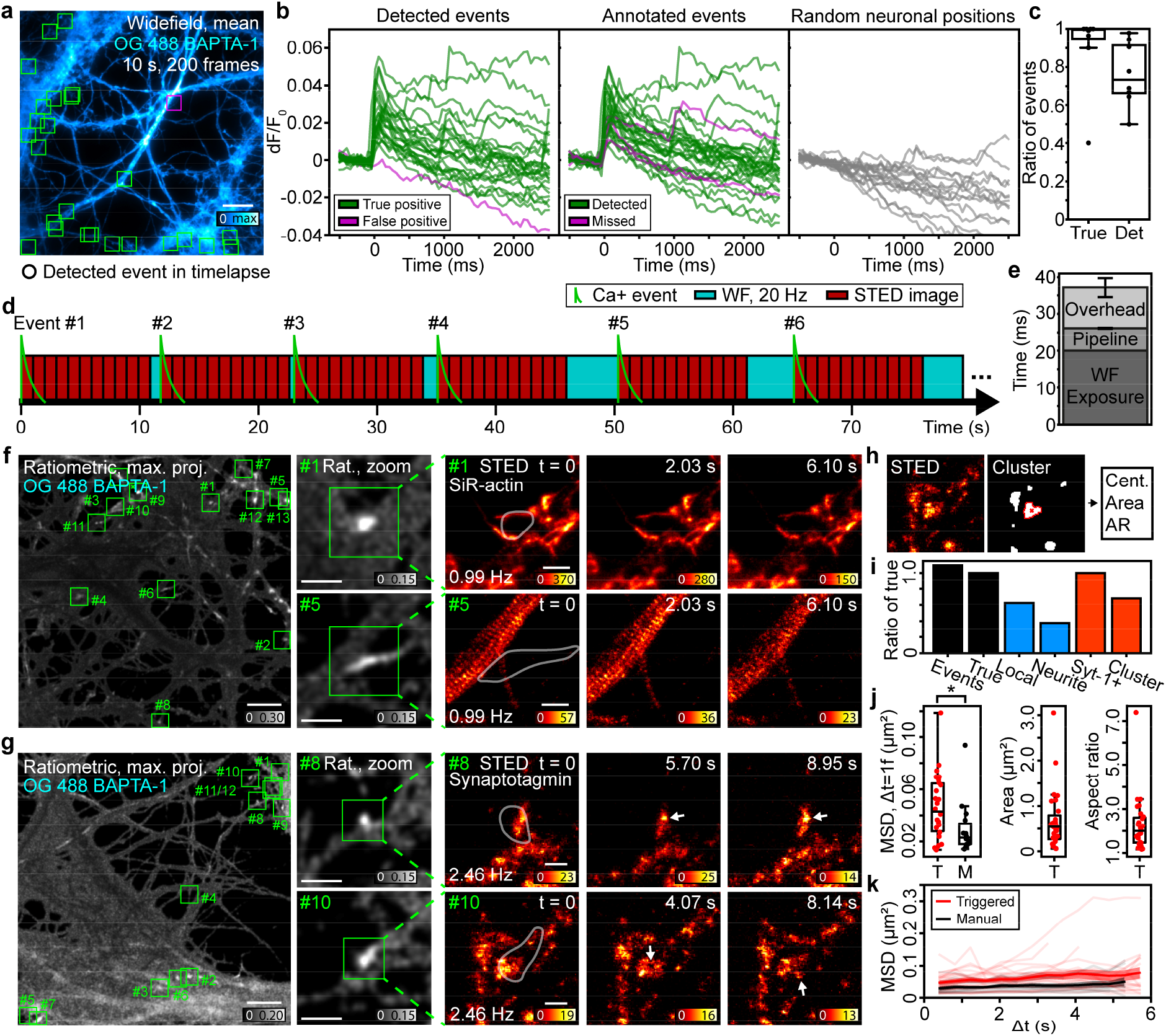
Neuronal functional and structural imaging with etSTED. **a**, Mean Oregon Green 488 BAPTA-1widefield image from a 20 Hz timelapse recording of 200 frames. Detected events are shown as boxes (green: true, magenta: false). N = 25 events. **b**, Extracted calcium curves in regions around the detected event coordinates (left, boxes marked in **a**), annotated event coordinates (center), or random neuronal positions (right). **c**, Characterization of the true positive detected event ratio (left) and detected annotated event ratio (right). N = 8 cells. **d**, Timeline of the experiment in **f**. **e**, Temporal performance of etSTED. N = 90 events in N = 11 cells. **f**,**g**, etSTED experiments with imaging of SiR-actin in **f** and Syt-1_STAR635P in **g**. Maximum-projected ratiometric image of all detected events in experiment (left, green squares show location of detected events), zoomed-in views of the ratiometric image in two detected events (center), and 0.99 Hz etSTED timelapse of the actin structures in **f**, and 2.46 Hz etSTED timelapse of syt-1_STAR635P in **g**. Overlayed semi-transparent white outline shows extent of detected local calcium activity. **h**, Analysis of synaptic vesicle clusters in active synapses. **i**, Event detection ratios, as compared to the number of true events, for number of total events (black, left), local calcium events (blue, left), neurite-wide calcium events (blue, right), etSTED timelapses with Syt-1-positive vesicles (orange, left), and etSTED timelapses with synaptic vesicle clusters (orange, right). N = 186 events in N = 17 cells. **j**, Analysis of synaptic vesicle clusters, in terms of mean square displacement (MSD) with Δt = 1 frame in calcium-triggered etSTED timelapses (red) and manual STED timelapses (black) (left), area (center), and aspect ratio (right). **k**, Analysis of the time-dependent MSD for synaptic vesicle clusters in calcium-triggered etSTED timelapses (red) and manual STED timelapses (black). Semi-transparent curves show individual clusters, solid lines show mean curves, and shaded areas show 83% confidence intervals. N = 26 clusters in N = 10 cells (red), N = 14 clusters in N = 14 cells (black). Scale bars, 10 μm (**a**,**f,g** ratiometric), 3 μm (**f**,**g** ratiometric zoom), 1 μm (**f** STED) and 500 nm (**g** STED). Box plots show 25–75% IQRs, middle line is mean, and whiskers reach 1.5 times beyond the first and third quartiles. Bar plots are means, and whiskers show 1 standard deviation.

A typical etSTED experiment in neurons can detect several calcium spikes sequentially and dynamically switches between the imaging techniques (Fig. 2d). The analysis of the temporal performance shows that, for a widefield imaging experiment with an exposure time of 20 ms and 800 x 800 pixels, the analysis pipeline takes 6.1 ± 0.4 ms to run (Fig. 2e). The analysis is implemented to run on a GPU, and thus the speed is improved tenfold over CPU-implementations (Supplementary Note 1). The time for the coordinate transform is < 1 ms. The overhead time, which includes toggling of hardware, scanning curve calculation, and scan initiation, is 11.0 ± 2.9 ms. Overall, the total triggering process from widefield exposure initiation to STED scan start in the case of a detected event, takes 37.1 ± 3.4 ms. This means that the maximum time lag between an event taking place and the STED scan starting at the site of that event is < 40 ms.

A maximum projected pre-processed ratiometric image over all events show clear areas of activity, both local calcium events, roughly < 5 μm^2^ in size, as well as spread-out neurite-wide events (Fig. 2f, left, middle). Upon triggering, STED imaging of actin (SiR-actin) was performed as a timelapse over 11 frames at 0.99 Hz in a 5 x 5 μm^2^ FOV (Fig. 2f, right, Supplementary Fig. 4). Three frames from two triggered STED timelapses show nanoscale actin structures in dendritic spines as well as the membrane periodic skeleton. The experiment lasted 2 min 46 s, from the detection of event 1 to the end of the STED timelapse following event 13 (Fig. 2f, green boxes). The duration of widefield imaging between finishing of the previous STED timelapse and the next detected event varied between 0–5 s (Fig 2d).

In the presynaptic compartment the voltage-dependent, local, and transient increase in intracellular calcium concentration leads, within few milliseconds, to synaptic vesicle exocytosis. ^15, 16^. The protein synaptotagmin-1 (Syt-1) is the calcium sensor that triggers vesicle release. While the dynamics of synaptic vesicles in the active zone has been studied^2^, their concurrent behaviour with the early stages of calcium enrichment is still unknown. We use etSTED to image the reorganization of synaptic vesicles in active synapses during Ca+ sensing (Fig. 2g, Supplementary Fig. 5, Supplementary Note 3). Antibodies against the intraluminal domain of Syt-1 are pre-mixed with secondary nanobodies conjugated to Abberior STAR635P and incubated with live neurons. Upon fusion of the vesicles with the plasma membrane, the Syt-1 intraluminal domain gets exposed to the extracellular milieu, allowing the binding of the nano-antibody^17^. When the synaptic vesicles are further recycled, the internalization of the labelled molecules enables imaging of the vesicle pools within the presynaptic active zone (Fig. 2g, right). The etSTED experiment had 12 detected Ca+ events and lasted 4 min 24 s (Fig. 2g, green boxes). We show ratiometric widefield images (Fig. 2g, middle) of two detected local events. The STED timelapses following the events (Fig. 2g, Supplementary Fig. 5), consisting of 31 frames taken at 2.46 Hz in a 3×3 μm^2^ FOV, show dynamic activity of the synaptic vesicles. In total, we imaged 186 events in 17 areas of cells, and 91% of the detected events were true calcium events (Fig. 2i, black), and out of those 62% were local calcium spikes and 38% were neurite-wide or larger events (Fig. 2i, blue). Furthermore, every true calcium spike event had Syt-1-positive vesicles in the vicinity, and 68% of them corresponded to clusters of synaptic vesicles (Fig. 2h–i, orange). Rearrangements of the shape, density, and connectivity of the densely packed synaptic vesicles such as disassembly of the clusters (Fig 2g, event #10) and individual vesicle dynamics (Fig 2g, event #8) are all captured in the STED timelapse recordings. We quantified the mean square displacement (MSD), area, and aspect ratio of the clusters in the active synapses (Fig. 2h). The area of the clusters was 0.68 ± 0.63 μm^2^ (Fig. 2j, middle), and the aspect ratio was 2.3 ± 1.2 (Fig. 2j, right). We also imaged similar vesicle clusters with manual STED imaging (Supplementary Note 4, Supplementary Fig. 6) and compared the MSD over the timelapses in the two cases (Fig. 2j, left, 2k). With calcium-triggered etSTED timelapse imaging we were able to record faster vesicle cluster dynamics which occur upon activity but especially within Ca+ oscillations. The MSD with Δt = 1 frame (0.41 s) was 0.046 ± 0.025 μm^2^ in the calcium-triggered timelapses and 0.030 ± 0.020 μm^2^ in the manual timelapses (two-sample two-sided Kolmogorov-Smirnov test: p = 0.018, test statistic: 0.48) (Fig. 2j, left). The time-dependent MSD furthermore shows a clear separation of the two curves at all Δt (Fig. 2k). Our method allows to monitor the synaptic vesicle dynamics during Ca+ oscillations and within the time domain of the kiss-and-run recycling mechanism, which resulted faster than those in averaged synapses without Ca+ synchronization.

In this work we outlined a new event-triggered method, etSTED, which enables super-resolved STED imaging inside < 40 ms from a detected event taking place, proving the potential for correlating millisecond dynamics such as calcium sensing in living cells with super-resolved details of proteins of interest. Imaging selected small regions around the detected event does not only minimizes cell stress and photodamage but also enables a significant increase of the temporal resolution in the regions of interest, in the order of 100–1000x faster as compared to manually acquiring STED timelapses in the whole investigated sample region. Moreover, it allows experiments not previously feasible, where the dynamics and structure of synaptic vesicle pools can be directly correlated to the presence or lack of local calcium oscillations. While the method as proposed here combines widefield imaging of a calcium sensor with STED imaging of actin, microtubules, or synaptic vesicles, the possibilities to generalize and extend the method are endless. Since the same control software could be used for any combination of imaging modalities, the super-resolved imaging can be extended to the likes of light sheet or RESOLFT microscopy, depending on the requirements on structure of interest, length of timelapse imaging, and spatial resolution. While our proposed peak detection pipeline is built for detecting instant intensity variations in the widefield images and proven to be efficient at localizing calcium activity and movement, detection of other fluorescent sensors, cellular processes, or specific movement could need tailored analysis pipelines to be accurately detected. The implementation in the control software allows for easy exchange to a pipeline optimized to the problem at hand and to the accuracy and speed of liking. The etSTED method bridges the gap between two types of imaging: small-scale, super-resolution imaging, and large-scale activity imaging, fostering new applications in functional and dynamic living systems.

## Material and Methods

### Microscopy set-up

All images were acquired on a custom-built and combined STED and widefield setup, based on a STED setup previously described^5^. A complete schematic and list of components can be found in Supplementary Fig. 2 and Supplementary Note 5. *STED*: Excitation of the red-shifted dyes was done with a pulsed diode laser at 640 nm with a pulse width of 60 ps (LDH-D-C-640, PicoQuant, Berlin, Germany). Depletion was done with a pulsed 775 nm laser beam with a pulse width of 530 ps (KATANA 08 HP, OneFive GmbH, Regensdorf, Switzerland). Fast on and off control of the excitation and depletion lasers for STED is done using an AOTF (AOTFnC-400.650-TN + MPDS4C-B66-22-74.156, AA Opto Electronic, Orsay, France) and AOM (MT110-B50A1.5-IR-Hk + MDS1C-B65-34-85.135-RS, AA Opto Electronic, Orsay, France) respectively, while for the widefield excitation laser it is done using the built-in fast modulation control. The depletion beam is shaped using a vortex phase mask on a spatial light modulator (LCOM-SLM X10468-02, Hamamatsu Photonics, Hamamatsu, Japan). A λ/4 and a λ/2 wave plate are used to create the circular polarization necessary for optimal depletion focus formation. Fast galvanometer mirrors are used for scanning in STED imaging (galvanometer mirrors 6215H + servo driver 71215HHJ 671, Cambridge Technology, Bedford, MA, USA) in a scanning system allowing constant resolution across an 80 x 80 μm^2^ FOV as described previously^5^. The fluorescence is decoupled with a dichroic mirror into the detection path. A bandpass filter (GT670/40m, Chroma Technology, Bellows Falls, VT, USA) and a notch filter (NF03-785E-25, Semrock Rochester, NY, USA) are used before the fluorescence is detected with an APD (SPCM-AQRH-13-TR, Excelitas Technologies, Waltham, MA, USA). *Widefield*: Excitation of Oregon Green 488 BAPTA-1 was done with a modulated continuous wave diode laser at 488 nm (Cobolt 06-MLD 488 nm, Cobolt, Solna, Sweden). Detection of the widefield images was done through a bandpass filter (FF01-540/80-25, Semrock) and a notch filter (ZET785NF, Chroma) using a sCMOS camera (Orca Flash 4.0, Hamamatsu Photonics). The widefield path is coupled into the beam path with a dichroic mirror after the scanning system. *General*: The setup uses a 100x/1.4 oil immersion objective (HC PL APO 100x/1.40 Oil STED White, 15506378, Leica Microsystems, Wetzlar, Germany) and a microscope stand (DMi8, Leica Microsystems). The system also uses a mechanical stage for moving of the sample in the lateral dimensions (SCAN IM 130 x 85 – 2 mm, Märzhäuser, Wetzlar, Germany) and a piezo to move the sample along the axial dimension (LT-Z-100, Piezoconcept, Lyon, France).

### Automatic control and PC

General control of the hardware in the microscope is performed using a NI-DAQ acquisition board (PCIe-6353, National Instruments, Austin, TX, USA). All hardware equipment is controlled with microscope control software ImSwitch^13^ written in Python. Control of the etSTED method is performed using a custom-written widget and controller in ImSwitch, available in GitHub (https://github.com/jonatanalvelid/ImSwitch-etSTED), that controls the lasers, image acquisition, and runs the real-time analysis pipeline with customizable parameters to adapt to sample-specific conditions.

Additionally, a focus lock that combines an infrared laser (CP980S, Thorlabs, Newton, NJ, USA), a CMOS camera (DMK 33UP1300, The Imaging Source Europe, Bremen, Germany), and the z-piezo through a feedback loop is coupled into the beam path just before the microscope stand. It is controlled with ImSwitch and allows for experiments to run without manual control for prolonged periods of time.

The PC used to control the complete microscope contains a Ryzen 7 3700X 8-core CPU (AMD, Santa Clara, CA, USA), and a GeForce RTX 3060 Ti TUF GAMING OC GPU (ASUS, Taipei, Taiwan).

### etSTED widget and optimized image analysis pipeline

The etSTED widget is built in a generalized way to allow the use of any combination of lasers and detectors controlled with ImSwitch. The widget allows any arbitrary analysis pipeline to be used, and it additionally contains a part for calibrating the coordinate transform between the image spaces of the two imaging modalities.

The coordinate transform utilised is a general third-order polynomial transformation that is optimized using Levenberg-Marquart optimisation. The transform is calibrated on the detected coordinates of the same objects in the two imaging spaces, for example fluorescent beads. The high-order polynomial allows it to be compatible and precise no matter what aberrations and distortions are present in the optical paths of the two modalities.

Pipelines are provided in the form of python functions and must use the current fast image as the input and returns a list of detected event coordinates in the fast-imaging space. The analysis pipeline used throughout this work is optimized to detect calcium events with Oregon Green 488 BAPTA-1 labelling in hippocampal neurons. It is schematically shown in Supplementary Figure 1, further described in Supplementary Note 1, and available in GitHub (https://github.com/jonatanalvelid/ImSwitch-etSTED). While it is optimized for hippocampal neurons, it also performs well in HeLa cells with the same labelling, and it is likely to perform well with similar and fast fluorescent sensor labels after tweaking of the pipeline parameters. The pipeline consists of two main parts: a pre-processing that uses the current frame, the previous frame, and a binary region-of-interest mask; and a peak detection. The pre-processing transforms the current image into a map of pixelwise percentual intensity changes as compared to the previous image. It does so by subtracting and dividing with the previous frame, as well as multiplying with the pre-calculated mask of the region of interest. After a final Gaussian smoothing the pre-processing outputs a map of the local changes in BAPTA-1 fluorescence, corresponding to the local changes in calcium level. The peak detection takes the pre-processed image and compares it with a maximum filtered version, where the coordinates of local maxima are found as the coordinates where the two are equal. After checking which of these are above a provided absolute ratiometric threshold by a mask multiplication, to avoid detection of fluorescence fluctuations due to noise, the coordinates of the detected events are sorted after ratiometric intensity and returned. In case of multiple detected peaks, the ratiometrically brightest peak is used throughout this work as the coordinate where the triggered STED imaging takes place. Most of the analysis pipeline (all matrix operations) runs on the GPU, significantly decreasing the runtime as compared to running it all on the CPU. Altogether, the optimized analysis pipeline runs in only 6 ms for the 800 x 800 pixels widefield images used in this work. Exact analysis pipeline parameters for the experiment of each supporting data acquisition can be found in Supplementary Table 1.

### Active synapses synaptotagmin-1 cluster analysis

The analysis of the larger synaptic vesicle clusters in each event-triggered etSTED timelapse and manual STED timelapse of active synapses was performed using the Fiji distribution of ImageJ and using the script provided, see Code Availability. To extract the clusters in each frame fairly, a histogram matching bleach correction step was firstly performed on the timelapses. Each frame was then smoothed with a 1-pixel Gaussian smoothing, and subsequently binarized with a global intensity threshold identical for each timelapse and 1-pixel eroded once. The binary objects larger than 0.015 μm^2^ were analysed for area, elliptical fitting for the aspect ratio, and centroid extraction. The data was then processed using Python and the scripts provided, where the traces of each cluster throughout the timelapses were connected using the centroids and a minimum-Euclidean-distance approach with an upper limit of 0.3 μm movement per frame. Then, the centroid traces of each cluster could be extracted, and for the largest cluster in the first frame in each timelapse, the trace was further analysed to extract the mean square displacement. The MSD was calculated for a specific Δ*t* as the mean of the Euclidean distance between each centroid position and that centroid position shifted with Δ*t* with respect to the first one throughout the timelapse.

### Primary neuronal culture

Primary neuronal cultures were prepared from embryonic day 18 (E18) Sprague Dawley rat embryos. The pregnant mothers were sacrificed with CO_2_ inhalation and aorta cut, and brains were extracted from the embryos. Hippocampi were dissected and mechanically dissociated in Minimum Essential Medium, MEM (Thermo Fisher Scientific, 21090022). 2 x 10^5^ cells per 60 mm culture dish were seeded on poly-D-ornithine (Sigma Aldrich, P8638) coated #1.5 18 mm glass coverslips (Marienfeld, 0117580), and were let to attach in MEM with 10% horse serum (Thermo Fisher Scientific, 26050088), 2 mM L-Glut (Thermo Fisher Scientific, 25030-024) and 1 mM sodium pyruvate (Thermo Fisher Scientific, 11360-070), at 37°C at an approximate humidity of 95–98% with 5% CO_2_. After 2–4 h, coverslips were flipped over an astroglial feeder layer (grown in MEM supplemented with 10% horse serum, 0.6% glucose, and 1% penicillin-streptomycin) and maintained in Neurobasal (Thermo Fisher Scientific, 21103-049) supplemented with 2% B-27 (Thermo Fisher Scientific, 17504-044), 2 mM L-glutamine and 1% penicillin– streptomycin. The neuronal cultures were treated with 5 μM 5-fluorodeoxyuridine (FDU) at DIV 2–3, to prevent glia overgrowth. The cultures were kept for up to 24 days and fed twice a week by replacing one-third of the medium per well: up to DIV7 with Neurobasal complete medium, and after DIV7 with BrainPhys (STEMCELL tech. 05790), 1% Pen/Strep (Gibco 15140-114) and SM1 Supplement (STEMCELL tech. 05711). The experiments were performed on mature cultures at DIV14–DIV21. All experiments were performed in accordance with animal welfare guidelines set forth by Karolinska Institutet and were approved by Stockholm North Ethical Evaluation Board for Animal Research. Rats were housed with food and water available ad libitum in a 12 h light/dark environment.

### HeLa culture

HeLa (ATCC CCL-2) cells were cultured in DMEM (Thermo Fisher Scientific, no. 41966029) supplemented with 10% (vol/vol) fetal bovine serum (Thermo Fisher Scientific, no. 10270106), 1% penicillin/streptomycin (Sigma-Aldrich, no. P4333) and maintained at 37 °C and 5% CO_2_ in a humidified incubator. Cells were plated on #1.5 18 mm glass coverslips (Marienfeld, 0117580) 24–48h before imaging.

### Active synapses labeling

For the labelling of active synapses, 1 μl of Synaptotagmin-1 antibody (1 mg/ml, luminal domain - Synaptic System, 105 3FB) and 1 μl of FluoTag-X2 anti-mouse KLC (batch n. N1020) conjugated to Abberrior STAR635P dye were pre-incubated with 98 μl of pre-conditioned neuronal medium, and incubated at room temperature (RT) for 20 min. Neurons were, then, incubated with the Synaptotagmin-1 labelling solution for 30 min in a humidified chamber at 37°C. After the incubation time neurons were left to recover for 5 min in their original medium and washed twice with artificial cerebrospinal fluid (ACSF) before the imaging. Imaging was performed in ACSF at RT.

### Actin and microtubules labeling

The labelling of F-actin and tubulin was performed, as previously described^18^, using Live Cell Fluorogenic F-actin Labelling Probe Kit (CY-SC001). 1 mM stock solution was obtained by dissolving 50 nmol SiR-actin or SiR-tubulin in 50 μL of anhydrous DMSO. For neurons, the stock solution was diluted 1:1000 in ACSF, for a final 1 μM staining solution. The neuronal culture was then incubated for 30 min at 37°C with the dissolved SiR-actin or SiR-tubulin, and washed twice in ACSF prior to the imaging. For HeLa cells, the stock solution was diluted 1:1000 in cell medium, for a final 1 μM staining solution. The cultured cells were then incubated for 30–45 min at 37°C with the dissolved SiR-actin or SiR-tubulin, and washed once in cell medium prior to imaging.

### Calcium imaging

For calcium imaging 10 ul of Pluronic acid F-127 solution (0.2 g of Pluronic acid in 1 ml of DMSO and shaken at 40°C for 20 min) was added to 50 μg of OregonGreen 488 BAPTA-1 AM (OG 488, Cat. No. 6257, BAPTA AM, Cat. No. 2787) for a 1 mM stock solution. The neuronal culture or HeLa culture was incubated for 30 min at 37°C with 1 μM OG 488 BAPTA-1 AM (1:1000 dilution in cell medium), and washed once in ACSF or cell medium prior to the imaging.

### Imaging conditions and acquisition parameters

Widefield images were recorded using 20 ms exposure time with a frame rate of 20 Hz, and in some cases with 50–100 ms exposure time with a frame rate of 10–20 Hz. The 488 nm laser power used was 0.30–0.35 mW for neurons and 1.9 mW for HeLa cells. STED images were recorded using a pixel size of 25–30 nm, dwell time of 30 μs, 640 nm laser power of 5–15 μW, and 775 nm laser power of 59–73 mW. Exact image acquisition parameters for the experiment of each presented image can be found in Supplementary Table 2. All laser powers were measured at the conjugate back focal plane of the objective lens, between the scan and the tube lens.

All presented images are raw data. For visualization purposes, all STED images have been smoothed with a 0.5-pixel Gaussian smoothing.

## Data availability

The data that support the implementation of the method and support the findings in this study, including images, log-files, and metadata, are openly available in Zenodo at https://doi.org/10.5281/zenodo.5593271, reference number 5593271.

## Author contributions

I.T. designed and supervised the research. J.A. and I.T. conceived the project idea. J.A. planned and built the setup; designed the control software and real-time analysis pipeline; planned, prepared, and performed the experiments; and performed the data analysis. M.D. prepared neuronal samples and provided guidance on neuronal experiments. J.A. and I.T. wrote the manuscript with input from all the authors.

## Code availability

The code of the control software and the specifically developed widget in ImSwitch used in this study is open source and available at https://github.com/jonatanalvelid/ImSwitch-etSTED. The code of the various analysis supporting the findings in this work, as ImageJ macros and Jupyter notebooks together with example data, are available at https://github.com/jonatanalvelid/etSTEDanalysis.

## Acknowledgements

I.T. thanks the Swedish Foundation for Strategic Research (project FFL15-0031) for supporting the research. F. Pennacchietti is acknowledged for help with the synaptotagmin image analysis.

## Notes

https://doi.org/10.5281/zenodo.5593271

### Competing Interest Statement

The authors have declared no competing interest.

