## Supplementary information for "Event-triggered STED imaging"

* Corresponding author: Ilaria Testa

0000-0003-4005-4997

**List of content**

1. **Real-time calcium activity detection pipeline**
2. **Post-acquisition true/false detected event decision**
3. **Example etSTED experiments**
4. **Manual STED timelapses of synaptotagmin-1**
5. **Microscope components**
6. **Supplementary figures**
7. **Supplementary tables**

**List of Figures**

1. Optimised calcium signal event detection analysis pipeline.
2. etSTED setup.
3. Post-acquisition true/false detected event decision.
4. etSTED experiment in neurons with calcium imaging (Oregon Green 488 BAPTA-1) and STED timelapse imaging of actin (SiR-actin).
5. etSTED experiment in neurons with calcium imaging (Oregon Green 488 BAPTA-1) and STED timelapse imaging of synaptic vesicles (synaptotagmin-1_STAR635P).
6. Manual STED timelapse imaging of synaptic vesicles (synaptotagmin-1_STAR635P).
7. etSTED experiment in neurons with calcium imaging (Oregon Green 488 BAPTA-1) and STED timelapse imaging of microtubules (SiR-tubulin).

### 1. Real-time calcium activity detection pipeline

The analysis pipeline used throughout the work is developed and optimised for detection of calcium activity spikes in hippocampal neurons in widefield images of Oregon Green 488 BAPTA-1 with a pixel size of 100 nm. The pipeline is used in the etSTED widget as a separate python function and uses the current and previous widefield images as input, and outputs the ratiometrically brightest detected event coordinates. The pipeline is schematically shown in Supplementary Fig. 2 and is available at https://github.com/jonatanalvelid/ImSwitch-etSTED/blob/master/smartsted/analysis_pipelines/bapta_calcium_spikes.py. The pipeline is developed in Python, mainly using the numpy[1], scipy[2], cupy, and opencv packages, and consists of two main parts: a pre-processing that uses the current frame, the previous frame, and a mask of the region of interest; and a peak detection algorithm, which detects any peaks in the resulting pre-processed image. Prior to initiating the etSTED method and running the pipeline, a binary mask representing the region to consider in the FOV, usually the cell, is generated. It is created by an intensity thresholding and Gaussian smoothing of a mean image of several consecutive frames, usually 10. This mask will be input to any detection pipeline applied, and limits any uncorrelated background noise, which contains large ratiometric changes from frame to frame, from affecting the result.

In the analysis pipeline, the initial pre-processing transforms the widefield image into a map of pixelwise percentual intensity change from the previous image. It does so by first subtracting the previous frame from the current and dividing with the previous frame, generating a ratiometric image of pixelwise percentual change in intensity. This image is then multiplied by the pre-calculated mask to only get the changes inside the region of interest and discard the background. The image is then subjected to a Gaussian smoothing step to lower the impact of noise-based fluctuations of the fluorescence intensity changes, and the final pre-processed image is an intensity-insensitive map of the local changes in BAPTA-1 fluorescence, corresponding to the local changes in calcium level. The intensity-insensitivity comes from looking at the ratiometric change of intensity rather than the raw intensity, and thus also small changes in calcium activity in dimmer local regions can be detected.

The subsequent peak detection uses a modified version of the peak detection algorithm used in *peak_local_max* in the scipy package. It takes the ratiometric image as input and compares it to a maximum filtered/dilated version of itself. The local maxima, and thus where we have intensity change peaks, are found as the coordinates where the two are equal. To avoid noise-based fluctuations of the fluorescent signal, such as Poisson and Gaussian noise, to affect our detection we multiply a Boolean mask with the intensity change peaks with a thresholded version of the ratiometric map. This absolute threshold on the intensity change ratio is perhaps the most important parameter of the analysis pipeline, and it must be adapted to the fluorescence signal level. Common values used in this work are between 0.05–0.15. Finally, peaks close to the border of the FOV are removed, to allow all the STED imaging to be performed inside the 80 x 80 $\mu m^2$ FOV. Then, the intensity values in the ratiometric image of the peaks that remain are extracted, and the brightest detected peaks are chosen, up to N peaks. The parameter N can be arbitrarily chosen in the widget and depends on how much information is wanted from the pipeline. While the triggered STED imaging will only be performed at the position of the ratiometrically brightest peak in case of multiple detected peaks, post-acquisition analysis of the data could benefit from knowing where other detected peaks in the same frame were located.

Prior to return of the coordinates, the coordinates can additionally be sorted to ensure a provided spatial spacing between each one of them, to avoid multiple detections of the same peak. This comes at the cost of calculation speed, and as the current implementation of etSTED only performs STED imaging at the ratiometrically brightest peak and minimization of the analysis pipeline speed is of highest interest, this is rarely useful. One could imagine implementations of etSTED where multiple detected peak sites are investigated simultaneously or subsequently with STED imaging, and there this sorting of final detected coordinates could be useful.

The modification of the peak detection from the implementation in scipy is allowing most of the pipeline to be run using the cupy package in Python and hence benefits from GPU-acceleration. For the size of widefield images used in this work, 800 x 800 pixels, this improves the runtime more than tenfold down to only 6 ms. This ultimately allows the method to start STED imaging inside < 40 ms from the point in time where any detected event occurs.

While the pipeline is designed and optimised to perform well in hippocampal neurons, it is also shown that it works well in HeLa cells. Moreover, the pipeline is most likely generalizable to imaging of other fluorescent sensors with rapid intensity changes (< 50 ms for 20 Hz imaging) by adapting the pipeline parameters accordingly.

### 2. Post-acquisition true/false detected event decision

To ensure that triggering events in the etSTED experiments performed throughout this work are true calcium signal events, a decision-making pipeline has been performed post-acquisition. During the acquisition, not only the last widefield frame and the STED images are saved, but also the N widefield frames leading up to the event, in our case N = 10, are saved along with a log file describing the triggering event (coordinates, timings, etc.). These frames can be used in post-acquisition analysis to determine whether the triggering event was in fact a true event or caused by something else than an increase in the calcium signal. As the analysis pipeline detected differences in the fluorescence signal from one frame to the next inside each pixel, fluorescent details moving in the image, as well as other causes of a fast fluctuation in the fluorescence signal, can also trigger the STED imaging.

To sort out the false events, the N widefield frames before the event can be investigated. By using the saved triggering coordinate in the log-file, the summed calcium signal trace surrounding the triggering coordinate, for example in a 5 x 5 $\mu m^2$ region, can be extracted up until the event was detected (Supplementary Fig. 3d). Any true calcium event will show a significant increase in the last frame, and any false event due to something moving fast in the frame to the detected pixel will show a relatively flat response. This is due to the fluorescence being present in surrounding pixels previously, assuming the object is not moving too fast for the 20 Hz widefield imaging to follow, and thus the movement is not causing any significant increase in the total fluorescence signal surrounding the affected pixel.

The real-time analysis pipeline could be improved performance-wise to not trigger on these events, by using the last few widefield frames instead of only the last, to analyse the calcium traces as described above. However, this as any other added real-time analysis would cause a significant decrease in speed of the total analysis pipeline, leading to slower times between the actual event and the triggered STED images, and it is thus a trade-off and something that could be adjusted depending on the application at hand. We choose to minimize the number and complexity of the steps of the pipeline and thus optimise the speed, as the falsely triggered events can be sorted out in post-acquisition analysis as shown here. In our experiments with neurons, moving filopodia are the most occurring false events, and we have not detected any false triggering events other than moving fluorescent objects that can be sorted out in the way described above.

### 3. Example etSTED experiments

We provide three examples of etSTED experiments with full STED timelapses and the widefield of each triggering event in Supplementary Fig. 4–6. Each experiment is performed in neurons, and the STED imaging is performed on microtubules (SiR-tubulin, Supplementary Fig. 7), actin (SiR-actin, Supplementary Fig. 4), or active synapses (synaptotagmin-1_STAR635P, Supplementary Fig. 5). All the recorded data in the experiments performed on these samples can further be found in the complementing shared dataset, see info under Data availability statement.

The experiment with etSTED imaging of microtubules (Supplementary Fig. 7) ran for 9 min 9 s and contained 16 detected events in total. Post-acquisition analysis determined 14 of them to be true events, and 2 false events caused by a moving filopodia. By investigating the triggering widefield frames and the calculated ratiometric images, 4 of the true events could be determined to be local calcium spikes in μm-sized limited regions such as single synapses, while the other 10 were found to be events with larger single or multiple neurite-wide signalling regions. The STED imaging was performed in regions of 5 x 5 $\mu m^2$ with a frame rate of 0.99 Hz for 30 frames. In the STED images we can see that the calcium events take place in regions and filaments rich of microtubule bundles. Thanks to the microtubules being present in almost all neurites and filaments, we can also confirm that the STED imaging takes place centered on the same area as the triggering event.

The experiment with etSTED imaging of actin (Supplementary Fig. 4) ran for 2 min 44 s and contained 12 detected events in total. Post-acquisition analysis determined 11 of them to be true events, and 1 false event caused by a moving filopodia. Of the true events, 4 could be determined to be local events, while the other 7 were found to be neurite-wide events. The STED imaging was performed in regions of 5 x 5 $\mu m^2$ with a frame rate of 0.99 Hz for 11 frames. In the STED images we see that the triggering calcium events happened in a mix of regions containing the membrane periodic skeleton, actin patches, and other structures.

The experiment with etSTED imaging of active synapses (Supplementary Fig. 5) ran for 3 min 9 s and contained 13 detected events in total. Post-acquisition analysis determined all of them to be true events. Of the true events, 9 could be determined to be local events, while the other 4 were found to be neurite-wide events. The STED imaging was performed in regions of 3x3 $\mu m^2$ with a frame rate of 2.46 Hz for 30 frames. In the STED timelapses we can follow the dynamics of the active synapses and presynaptic vesicles. In all events, we see active rearrangement of the clusters of presynaptic vesicles and movement of individual synaptic vesicles.

### 4. Manual STED timelapses of synaptotagmin-1

To compare the dynamics of the clusters of presynaptic vesicles in active synapses detected in the calcium-triggered STED images (Fig. 2g, Supplementary Fig. 5) to the dynamics without the presence of a calcium signal, we manually imaged STED timelapses of regions with the same size and frame rate and in the same samples, without considering the calcium signals

(Supplementary Fig. 6a). The regions to image were chosen as regions with synaptotagmin-1 clusters in a confocal scan of the full 80 x 80 $\mu m^2$ field of view. To ensure that we were looking at similar clusters of presynaptic vesicles, we measured the area and aspect ratio of the largest cluster in each region of interest. The areas and aspect ratios of the clusters imaged in this way was not different to that of the clusters in the etSTED timelapses (Supplementary Fig. 6b). The mean area was 0.68 ± 0.63 $\mu m^2$ for the clusters in the calcium-triggered events, and 0.63 ± 0.52 $\mu m^2$ for the clusters in the manual timelapses. The aspect ratio was 2.3 ± 1.2 for the clusters in the calcium-triggered events, and 2.1 ± 0.7 for the clusters in the manual timelapses. Two-sided two-sample Kolmogorov-Smirnov tests performed on the distributions returned p = 0.90 (test statistic = 0.17) for the area, and p = 0.76 (test statistic = 0.20) for the aspect ratio.

### 5. Microscope components

The following is a list of the microscope components as labelled in Supplementary Fig. 2.

**Lenses**: L1: 300 mm, L2, L4, L15: 200 mm, L3: 100 mm, L5: 400 mm, L13: 150 mm (all AC254-XXX-B-ML, Thorlabs), L6, L9: 100 mm, L7, L8: 200 mm, L10: 75 mm, L14: 30 mm (all AC254-XXX-A-ML, Thorlabs), L11: 250 mm, L12: 150 mm (both AC508-XXX-A-ML, Thorlabs), SL: 50 mm (Leica), TL: 200 mm (Leica), OBJ: HC PL APO 100x/1.40 Oil STED White (15506378, Leica). **Filters**: BPF1: ET705/100m (Chroma), BPF2: FF01-540/80 (Semrock), NF1: NF03-785E (Semrock), NF2 and NF3: ZET785NF (Chroma), CUF: CT780/20bp (Chroma). **Dichroic mirrors**: DM1: T700dcspxruv_UF3 (Chroma), DM2: ZT405/488/561/640/775rpc (Chroma), DM3: T860SPXRXT (Chroma), DM4: FF552-Di02 (Semrock), DM5: Di02-R488 (Semrock). **Lasers**: 488: 06-MLD 488 nm (488 nm, Cobolt), 640: LDH-D-C-640 (640 nm, PicoQuant), 775: KATANA 08 HP (775 nm, OneFive), 980: CP980S (Thorlabs). **Detectors/cameras**: APD: SPCM-AQRH-13-TR (Excelitas), WF-CMOS: ORCA-Flash4.0 v2 (Hamamatsu), FL-CMOS: DMK 33UP1300 (The Imaging Source Europe GmbH, Bremen, Germany). **Scanners**: GX/GY: 6215H Galvanometric mirrors + 71215HHJ 671 Servo Driver (Cambridge Technology), Z-piezo: Z-piezo stage LT-Z-100 (Piezoconcept, Lyon, France), XY-stage: SCAN IM 130 x 85 – 2 mm (Märzhäuser Wetzlar GmbH, Wetzlar, Germany). **Fiber optics**: F1: PMJ-3AHPM3S-633-4/125-3-3-1 (OZ Optics), F2, F3: P5-488PM-FC-2 (Thorlabs), FC1, FC2: 60SMS-1-0-A2-02, FC3: 60SMS-1-0-A18-02, FC4: 60FC-4-M4-33 (all Schäfter + Kirchhoff GmbH, Hamburg, Germany). **Misc.**: PH: P75H (Thorlabs), AP: D15S (Thorlabs), PBS: PTW 1.15 (Bernard Halle Nachfl. GmbH, Berlin, Germany), λ/4-1: 600-1200 achr. (RAC 5.4.15, B.Halle Nachfl.), λ/4-2: 460-680 achr. (RAC 3.4.15, B.Halle Nachfl.), λ/4-3 and λ/4-4: 500-900 achr. (RAC 4.4.15, B.Halle Nachfl.), λ/2: 500-900 achr. (RAC 4.2.15, B.Halle Nachfl.), SM: 50 mm concave mirror (CM508-050-P01, Thorlabs), SLM: LCOS-SLM X10468-02 (Hamamatsu Photonics), AOM: MT110-B50A1.5-IR-Hk + MDS1C-B65-34-85.135-RS (AA Opto Electronic), PSDB: PSD-065-A-MOD (Micro Photon Devices, Bolzano, Italy), Mirrors: BB1-E02 and PF10-03-P01 (Thorlabs). **PC:** CPU: Ryzen 7 3700X 8-core (AMD), GPU: GeForce RTX 3060 Ti TUF GAMING OC (ASUS).

### 6. Supplementary figures

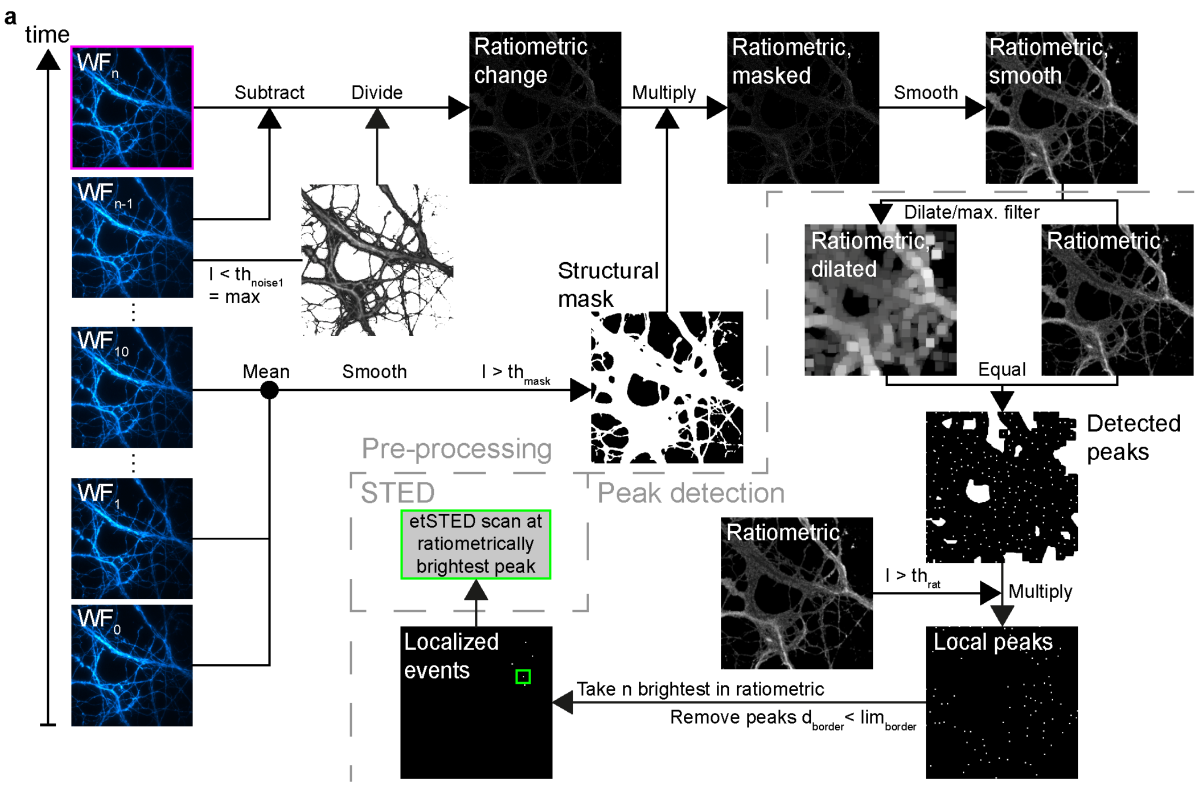


**Supplementary Figure 1. Optimised calcium signal event detection analysis pipeline. a,** Schematic view of the analysis pipeline used to detect calcium signal events with BAPTA-1-labelled cells.

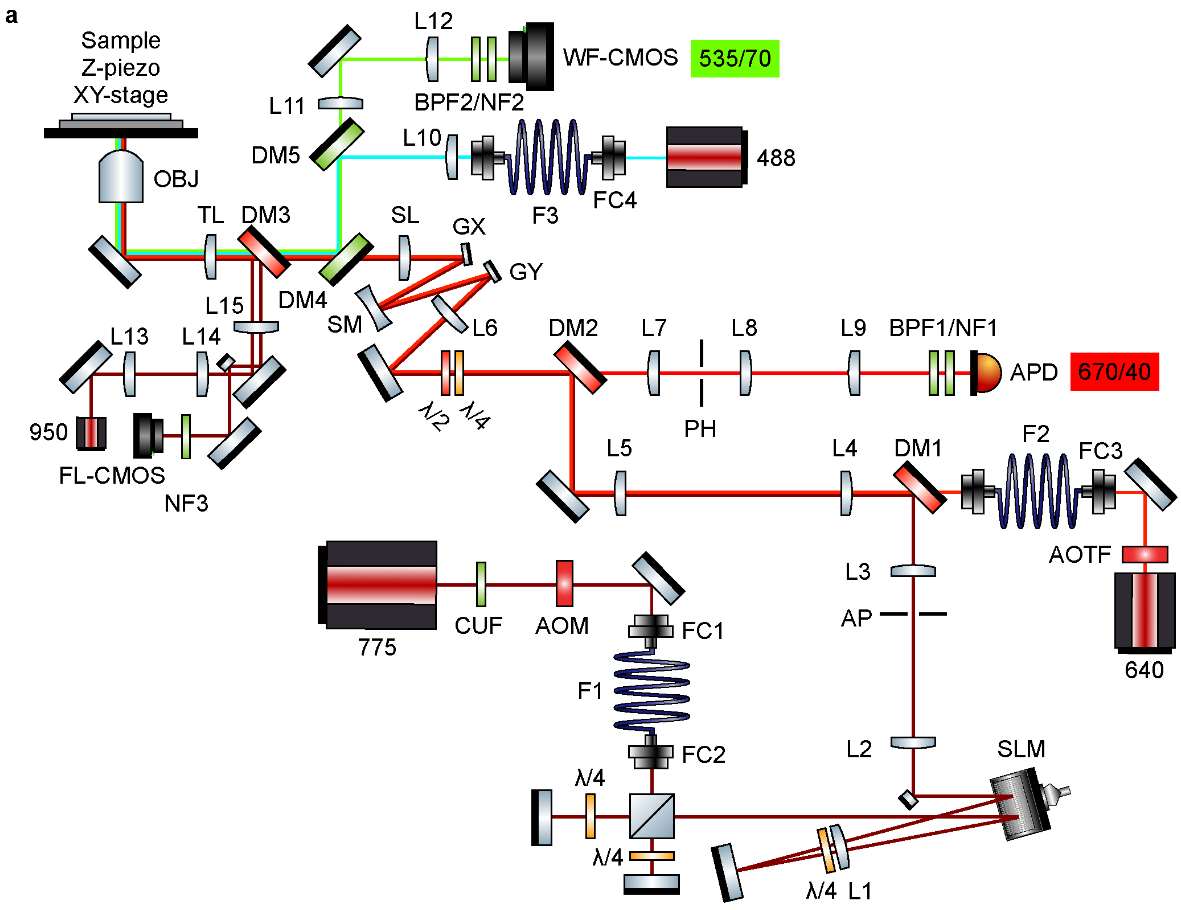


**Supplementary Figure 2. etSTED setup. a**, Extended schematic view of the etSTED setup, including STED, widefield, focus lock, and sample modules.

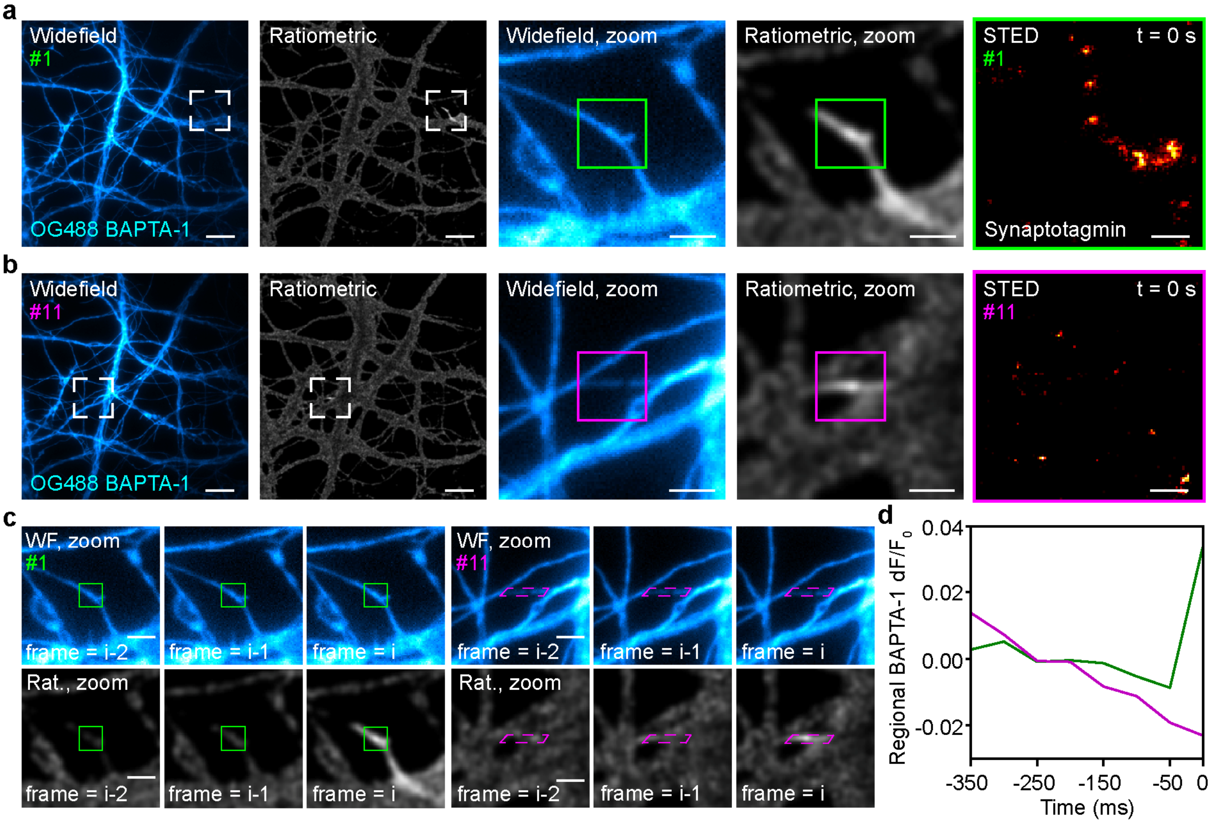


**Supplementary Figure 3. Post-acquisition true/false detected event decision. a**, Example widefield, ratiometric, and STED image from a true detected calcium event. White dashed boxes mark the zoomed-in regions (widefield, ratiometric). Green boxes mark the etSTED-imaged region (widefield, zoom; ratiometric, zoom). **b**, Example widefield, ratiometric, and STED image from a false detected calcium event. White dashed boxes mark the zoomed-in regions (widefield, ratiometric). Magenta boxes mark the etSTED-imaged region (widefield, zoom; ratiometric, zoom). **c**, Zoom-in of the widefield (top) and ratiometric (bottom) images for the three time points (frame = i-2, i-1, i) leading up to the detected event (frame = i), in the true (left) and false (right) event. Green boxes mark a region with increased calcium signal (true event). Magenta boxes mark a region with a moving filament (false event). **d**, Regional Oregon Green 488 BAPTA-1 fluorescence difference signal in a 5 x 5 μm$^2$ region around the detected event coordinate before the triggering event and in the triggering frame. Scale bars, 10 μm (**a**,**b** widefield, ratiometric), 2 μm (**a**,**b** zooms, **c**) and 500 nm (**a**,**b** STED).

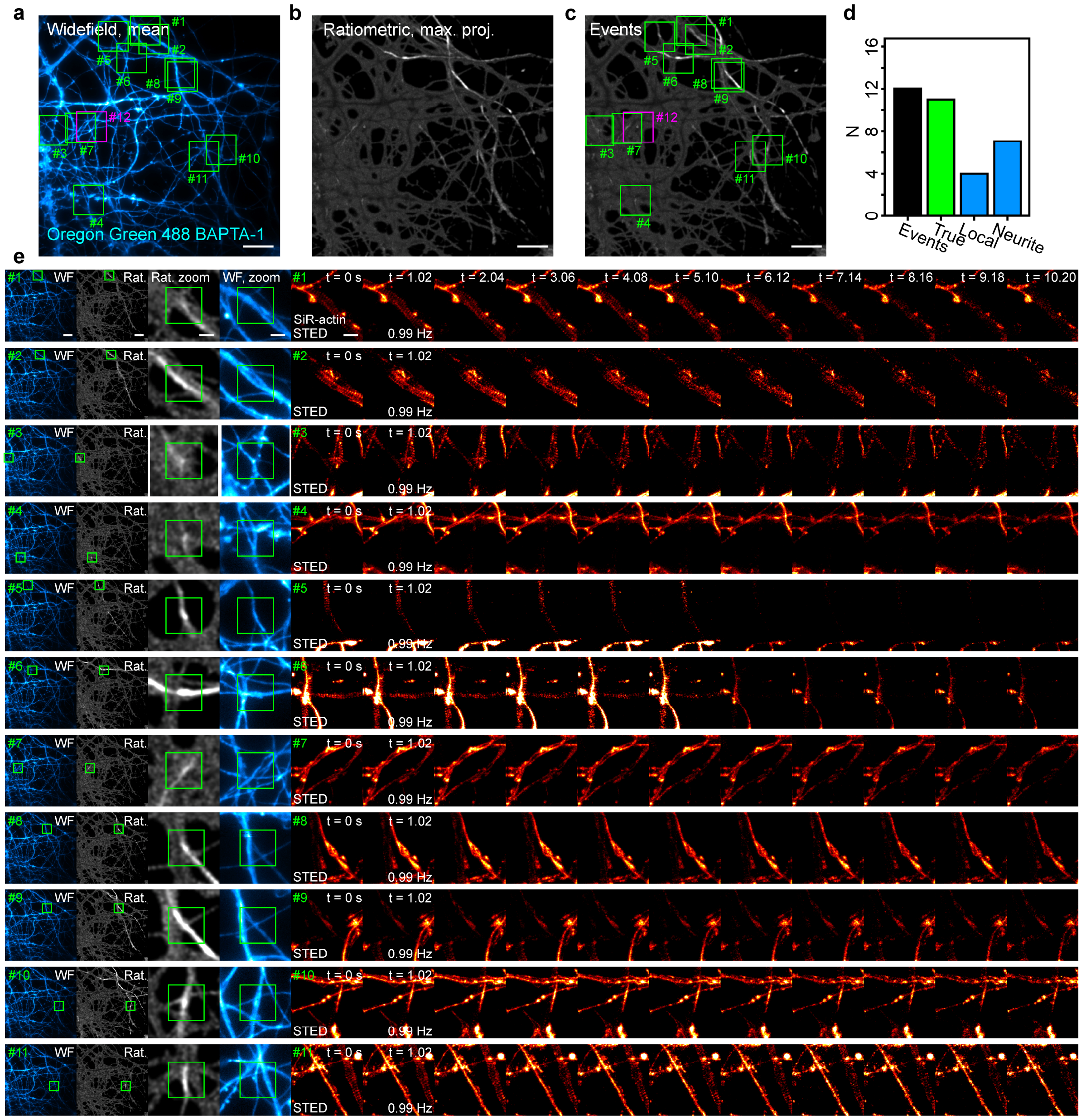


**Supplementary Figure 4. etSTED experiment in neurons with calcium imaging (Oregon Green 488 BAPTA-1) and STED timelapse imaging of actin (SiR-actin). a**, Mean image of all widefield frames with detected events. Boxes are centered on the coordinates of true (green) and false (magenta) detected events. **b**, Maximum projection of all ratiometric preprocessed widefield frames with detected events. **c**, Same as **b** with boxes centered on the coordinates of true (green) and false (magenta) detected events. **d**, Number of detected events, true events, local events, and neurite-wide events. **e**, Widefield frame, ratiometric preprocessed frame, zoom-ins of the widefield and ratiometric, and event-triggered STED timelapse (11 frames, 0.99 Hz) of the events as numbered and marked in **a**,**c**. Boxes marks the center of the detected event. Same scales and time labels apply to all timelapses. Scale bars, 10 µm (**a**,**b**,**c**,**e** widefield and ratiometric), 2 µm (**e** zooms) and 1 µm (**e** STED).

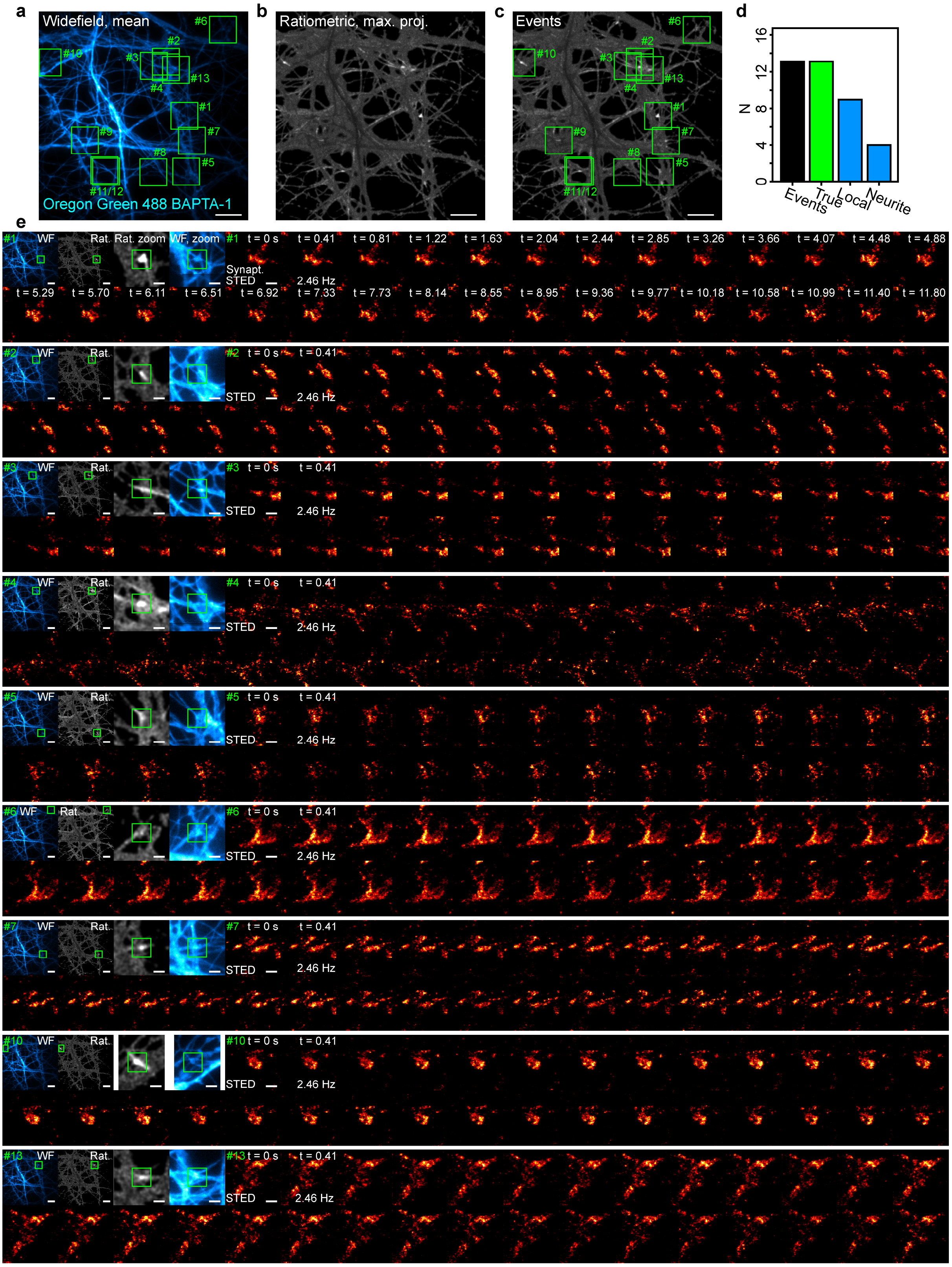


**Supplementary Figure 5. etSTED experiment in neurons with calcium imaging (Oregon Green 488 BAPTA-1) and STED timelapse imaging of synaptic vesicles (synaptotagmin-1_STAR635P). a**, Mean image of all widefield frames with detected events. Boxes are centered on the coordinates of true (green) and false (magenta) detected events. **b**, Maximum projection of all ratiometric preprocessed widefield frames with detected events. **c**, Same as **b** with boxes centered on the coordinates of true (green) and false (magenta) detected events. **d**, Number of detected events, true events, local events, and neurite-wide events. **e**, Widefield frame, ratiometric preprocessed frame, zoom-ins of the widefield and ratiometric, and event-triggered STED timelapse (30 frames, 0.99 Hz) of the events as numbered and marked in **a**,**c**. Boxes marks the center of the detected event. Same scales and time labels apply to all timelapses. Scale bars, 10 μm (**a**,**b**,**c**,**e** widefield and ratiometric), 2 μm (**e** zooms) and 1 μm (**e** STED).

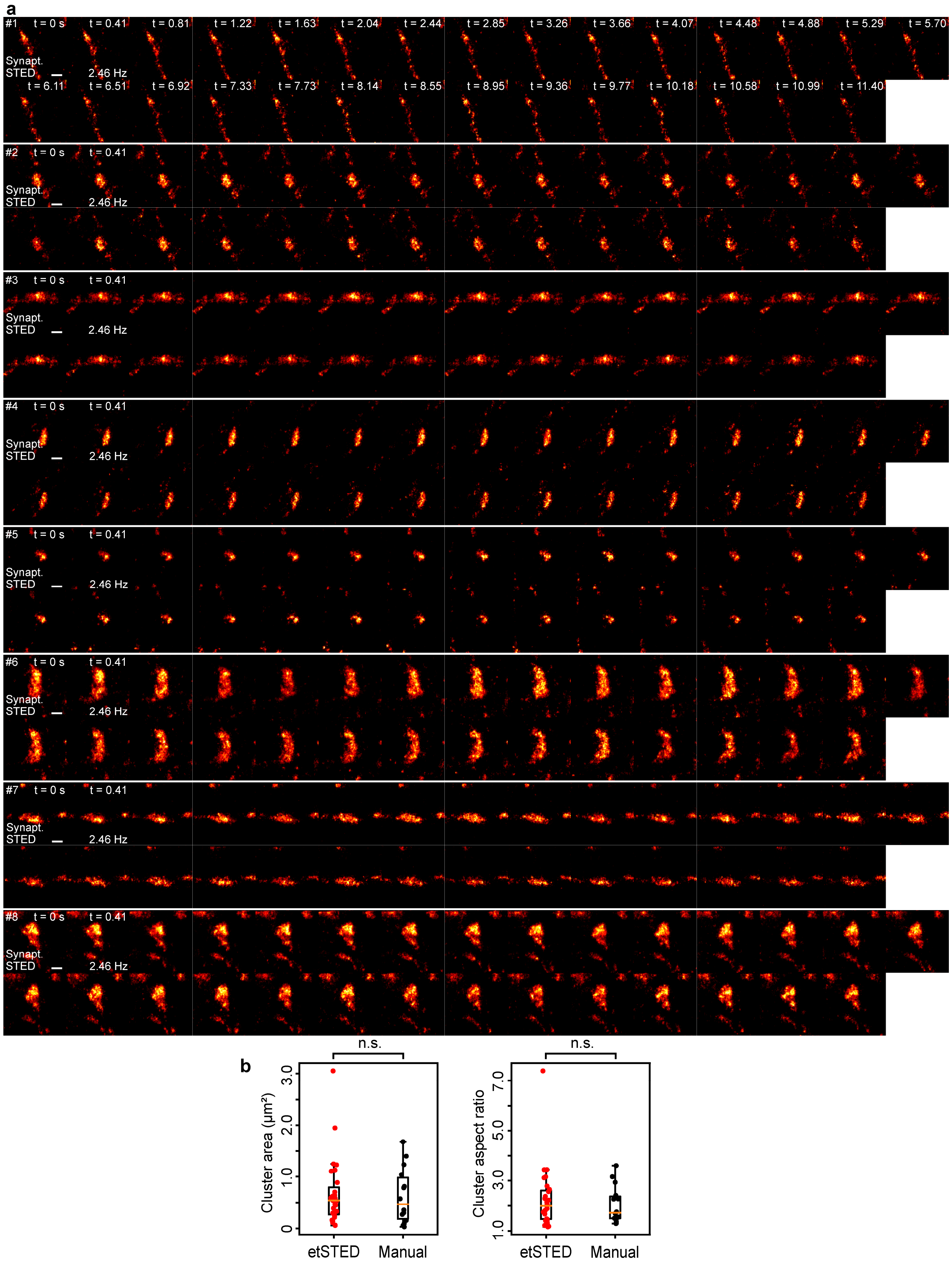


**Supplementary Figure 6. Manual STED timelapse imaging of synaptic vesicles (synaptotagmin-1_STAR635P). a**, STED timelapses of presynaptic vesicle clusters in active synapses. **b**, Distributions of area and aspect ratio for the clusters in calcium-activity-triggered STED timelapses (red) and manual timelapses (black).

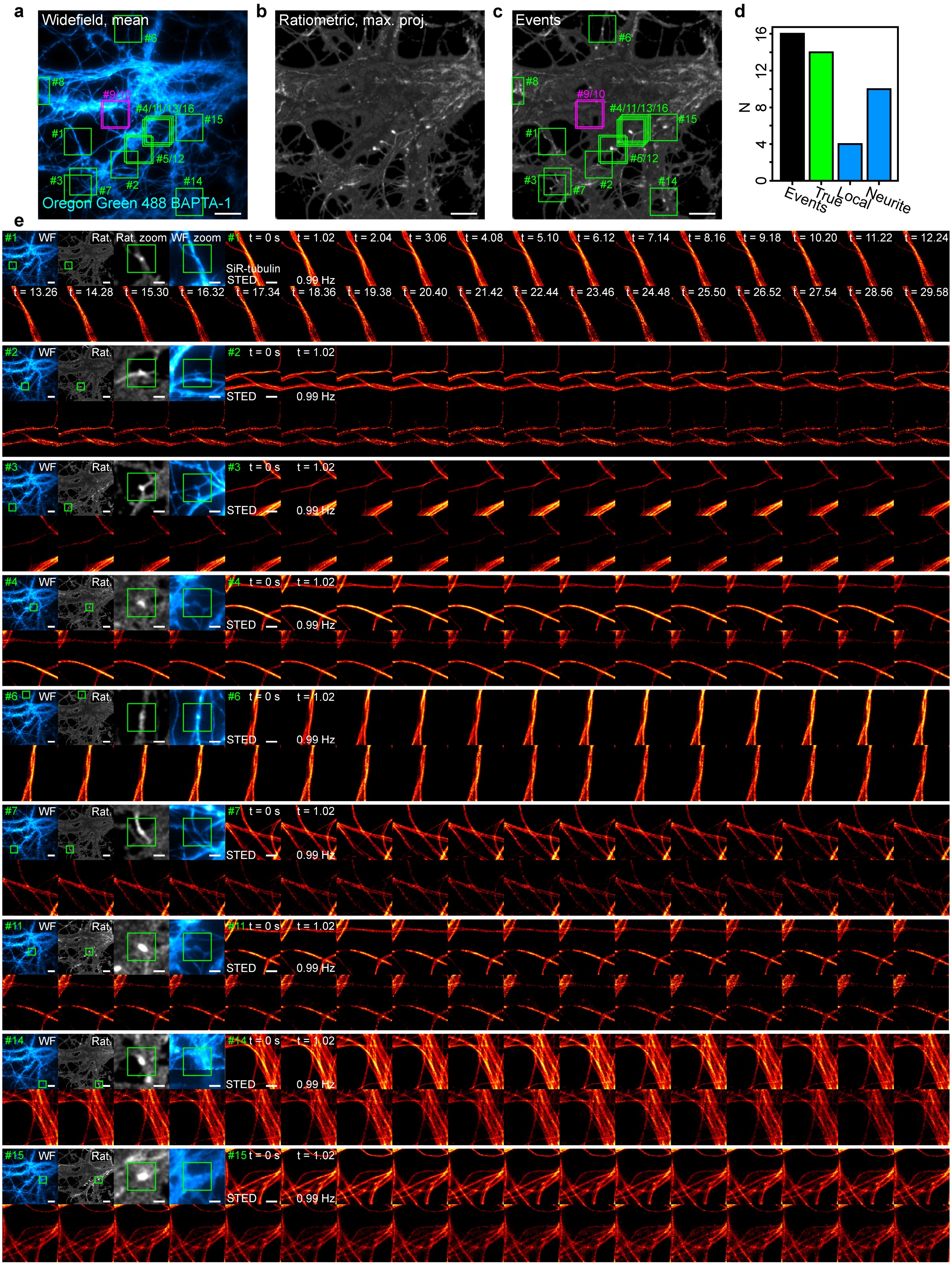


**Supplementary Figure 7. etSTED experiment in neurons with calcium imaging (Oregon Green 488 BAPTA-1) and STED timelapse imaging of microtubules (SiR-tubulin). a**, Mean image of all widefield frames with detected events. Boxes are centered on the coordinates of true (green) and false (magenta) detected events. **b**, Maximum projection of all ratiometric preprocessed widefield frames with detected events. **c**, Same as **b** with boxes centered on the coordinates of true (green) and false (magenta) detected events. **d**, Number of detected events, true events, local events, and neurite-wide events. **e**, Widefield frame, ratiometric preprocessed frame, zoom-ins of the widefield and ratiometric, and event-triggered STED timelapse (30 frames, 0.99 Hz) of the events as numbered and marked in **a**,**c**. Boxes marks the center of the detected event. Same scales and time labels apply to all timelapses. Scale bars, 10 µm (**a**,**b**,**c**,**e** widefield and ratiometric), 2 µm (**e** zooms) and 1 µm (**e** STED).

### 7. Supplementary tables

**Supplementary Table 1.** etSTED analysis pipeline parameters for the data presented in the figures.

| Fig | min_dist | thresh_abs | num_peaks | noise_level | smoothing_radius | ensure_spacing | border_limit |
|---|---|---|---|---|---|---|---|
| **1a** | 20 | 0.12 | 5 | 250 | 2 | 0 | 10 |
| **1c** | 20 | 0.06 | 5 | 700 | 2 | 0 | 10 |
| **2a** | 20 | 0.08 | 5 | 350 | 2 | 0 | 10 |
| **2f** | 20 | 0.12 | 5 | 500 | 2 | 0 | 10 |
| **2g** | 20 | 0.12 | 5 | 220 | 2 | 0 | 10 |
| **S3** | 20 | 0.12 | 5 | 220 | 2 | 0 | 10 |
| **S4** | 20 | 0.12 | 5 | 250 | 2 | 0 | 10 |
| **S5** | 20 | 0.10 | 5 | 500 | 2 | 0 | 10 |
| **S6** | 20 | 0.12 | 5 | 220 | 2 | 0 | 10 |

**Supplementary Table 2.** etSTED sample and image acquisition parameters for widefield and STED for the data presented in the figures.

| **General** | | | **Widefield** | | | | | **STED** | | | | | | |
|---|---|---|---|---|---|---|---|---|---|---|---|---|---|---|
| **Fig** | **Sample** | **Labels** | **$P_{488}$ (mW)** | **Img size (µm)** | **Px size (nm)** | **Exp time (ms)** | **Frame rate (Hz)** | **$P_{640}$ (µW)** | **$P_{775}$ (mW)** | **Img size (µm)** | **Px size (nm)** | **Dwell time (µs)** | **Frame rate (Hz)** | **Frames** |
| **1a** | Neurons | OG 488 BAPTA-1, SiR-tubulin | 0.31 | 80 | 100 | 20 | 20 | 5.0 | 73 | 3 | 30 | 30 | - | - |
| **1c** | HeLa | OG 488 BAPTA-1, SiR-tubulin | 1.90 | 80 | 100 | 100 | 10 | 8.5 | 61 | 2 | 25 | 30 | - | - |
| **2a** | Neurons | OG 488 BAPTA-1 | 0.35 | 80 | 100 | 50 | 20 | - | - | - | - | - | - | - |
| **2f** | Neurons | OG 488 BAPTA-1, SiR-actin | 0.35 | 80 | 100 | 50 | 20 | 15 | 60 | 5 | 30 | 30 | 0.99 | 11 |
| **2g** | Neurons | OG 488 BAPTA-1, Synaptotagmin-1_STAR635P | 0.31 | 80 | 100 | 20 | 20 | 12 | 59 | 3 | 30 | 30 | 2.46 | 31 |
| **S3** | Neurons | OG 488 BAPTA-1, Synaptotagmin-1_STAR635P | 0.31 | 80 | 100 | 20 | 20 | 12 | 59 | 3 | 30 | 30 | 2.46 | 31 |
| **S4** | Neurons | OG 488 BAPTA-1, SiR-tubulin | 0.30 | 80 | 100 | 20 | 20 | 5.0 | 73 | 5 | 30 | 30 | 0.99 | 31 |
| **S5** | Neurons | OG 488 BAPTA-1, SiR-actin | 0.35 | 80 | 100 | 20 | 20 | 15 | 60 | 5 | 30 | 30 | 0.99 | 11 |
| **S6** | Neurons | OG 488 BAPTA-1, Synaptotagmin-1_STAR635P | 0.31 | 80 | 100 | 20 | 20 | 12 | 59 | 3 | 30 | 30 | 2.46 | 31 |
| **S7** | Neurons | Synaptotagmin-1_STAR635P | - | - | - | - | - | 12 | 59 | 3 | 30 | 30 | 2.46 | 31 |
